# The Pleistocene-Holocene aquatic molluscs as indicators of the past ecosystem changes in Transbaikalia (Eastern Siberia, Russia)

**DOI:** 10.1101/2020.06.19.161216

**Authors:** Olga K. Klishko, Evgeniy V. Kovychev, Maxim V. Vinarski, Arthur E. Bogan, Georgiy. A. Jurgenson

## Abstract

Data on historical change of the Transbaikalian malacofauna in the Neopleistocene and Holocene is presented. Fossil shells from archeological excavations of the ancient settlements dating from the Neolithic period to Medieval and also from a drill hole of the Neopleistocene alluvial deposits were collected. In total nine species of bivalve molluscs from the families Margaritiferidae, Unionidae, Limnocardiidae, Glycymerididae, including one marine species, and two gastropod species from families Viviparidae and Planorbidae were identified. The time of the existence of each fossil species was determined by radiocarbon dating. It was found that the species ranged in age from more 50,000 and 2,080–1,210 years ago. Five species inhabited the Transbaikal region and are locally extirpated in the present. Their disjunctive ranges in the past included southern Europe and Western and Eastern Siberia to Transbaikalia and in the east to Far East and Primorye of Russia. The time of existence and extirpation of the thermophilic species of genera *Adacna, Planorbis, Lanceolaria* and *Amuropaludina* corresponds to cycles of the warming and cooling in Pleistocene and Holocene according to regional climate chronological scales. It was possible to separate these species as indicators of paleoclimate. Change of the species composition of the malacofauna of region connected with natural cycles of climatochrons in the Pleistocene and Holocene is the appearance of the climatogenic succession. In the course of this succession the disappearance of the stenothermal species occurred on a regional level and decreasing their global ranges.

## INTRODUCTION

Freshwater molluscs, both gastropods (snails) and bivalves (clams, mussels), represent a very significant component of freshwater ecosystems throughout the world [1], [2], [3]. This animal group is characterized by high species richness (more than 6000 valid species in the modern fauna [4], ecological plasticity and broad distribution. Freshwater molluscs can be found in all continents, except Antarctica, and dwell in habitats of different types, from large rivers and lakes (Lake Baikal) to ephemeral ponds, underground springs or rivers, and thermal pools. It is an ancient group of invertebrates, whose geological record can be traced back to the mid-Paleozoic [5], [6]. Since the mid-19^th^ century, shells of freshwater Mollusca are recognized as an important source of data for knowledge of the past ecosystems, their taxonomic diversity, physical and chemical characteristics (depth, temperature, salinity, and other indicators). Mollusc shells are easily fossilized by virtue of their being robust in composition, and, in many cases, there is a possibility to obtain large samples of mollusc remains from different rock horizons that allows one to reconstruct both phylogeny and patterns of their evolution. The usefulness of paleomalacological studies for reconstructions of the climate and landscape dynamics has been repeatedly stressed in the literature [7], [8], [9], [10], [11]. Some taxonomic groups of bivalves (like the Unionidae, a diverse family with more than 840 recent species) are recognized as good indicators of the temperature regime and climate changes as well as the anthropogenic impact [12], [13], [14], [15], [16], [17], [18]. Historical shifts in distribution of particular species of freshwater Mollusca reflect past alterations in landscapes and connections of paleobasins. The data on the past occurrences of mollusc species may be helpful for modern conservation efforts [19].

Traditionally, most information on fossil molluscs of the Pleistocene and the Holocene is obtained in the course of geological and paleontological investigations. However, there is another important source of primary data, namely, archeological excavations, which often provide numerous remains of shells. The modes of exploitition of molluscs by prehistoric people are numerous and range from eating them to using their shells for ornamental purposes or the manufacture of tools [20], [21], [22], [23]. To the best of our knowledge, the use of mollusc shells found during archeological excavations for paleoecological reconstructions is still rather limited (at least as compared to analogous studies carried out by paleogeographers and paleontologists).

The main goal of this study was to use the freshwater shells from long-term archeological excavations and a borehole made in the Transbaikalia area (Eastern Siberia, Russia) for a better understanding of the process of the formation of the regional malacofauna in context of the past climate changes.

Transbaikalia is characterized by a sharp and frequent spatial and temporal variability of climate, associated with the mountainous nature of the relief, the basin effect, vertical zonality, etc. [24], [25], [26], [27]. In the regional climatic chronological scale of Transbaikalia, several climatic rhythms of cryostages and thermal stages at the turn of 50–10 thousand years ago (Paleolithic) and 10–1 thousand years ago (Mesolithic-Iron Age period) are recognized [27]. The alternation of periods of warming and cooling in the Pleistocene (300-10 thousand years ago) and Holocene (10-1 thousand years ago) in the north of Transbaikalia is clearly recorded by changing the spore-pollen complexes of plant communities, which are the most sensitive to the dynamics of environmental factors [28], [29], [30], [31].

The aims of our work follows. 1. To determine both taxonomic identity and the absolute age of freshwater shell remains from Transbaikalia; 2. To reveal mollusc species able to serve as the climate change indicators; 3. To correlate the changes in the malacofauna species composition with the climatic rhythms in the Neopleistocene and Holocene of Transbaikalia.

### MATERIAL AND METHODS

We examined the extensive conchological material collected in 1975–2019 during archeological excavations in the territory of Transbaikalia [32], [33], [34], [35]. These excavations have been carried out in archaeological sites, remains of settlements, and the burial grounds of prehistoric humans living in the Transbaikalia area (Fig. 1A). Some shells collected in 2019 from a drill-hole located in the Upper Amur Basin were also used. Most excavation were made under supervision of E.V. Kovychev and included the systematic study of archeological objects and recovered artefacts. All investigated objects belonged to archeological sites dated from the Stone Age to Medieval (11–3 – 1.2–0.2 thousand years ago). Excavations were carried out on stratigraphy blocks at depths between 0.8 and 6.0 m. Each occupation layer amounted to 10–15 cm. All finds (bones of wild and domestic animals, ceramics, bitch bark, crafts made of bone and metal, fragments and shells of molluscs) were sorted, classified, described, inventoried and dated.

**Fig. 1.**
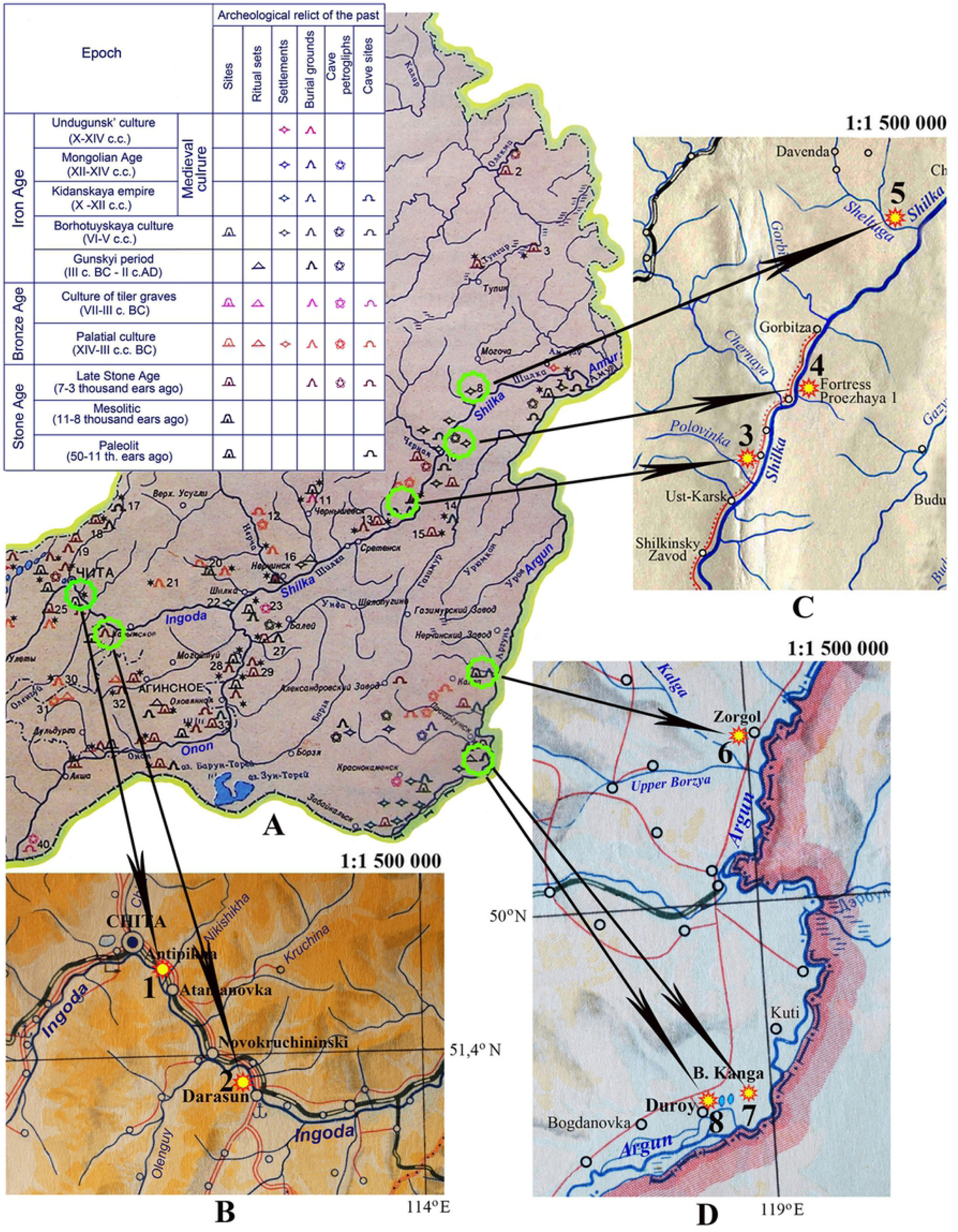
Collection sites of malacological material. **A** – archeological map of Transbaikalia [34] by circles marked the archeological old relict, by numerals – collection sites of fossil shells. **B1** – Chitinskyi region, Antipikha settlement, Ingoda River high-water bed, drill hole (Neopleistocene); **B2** – Karymsk’s region, Darasun settlement, ancient site. **C** – Sretensk’s region: **3** – ancient Luzhanky settlement (late Stone Age), **4 –** fortress Proezzhaya 1 (X-XII century AD), **5** – ancient settlement by Zheltuga River (late Stone Age). **D** – Priargunsk’s region: **6** – burial Zorgol (late Stone Age), **7** – Bol’shaya Kanga ancient settlement 2 (late Stone Age); **8** – burial Duroy 6 (late Stone Age).

The absolute age of the fossil shells was determined by radiocarbon dating performed at the Institute of the Earth Sciences of the Saint-Petersburg State University, in the laboratory “Geomorphological and paleontological investigations of the Polar regions and the World ocean”. The calibrated (calendar) age was determined by means of a calibrating software «OxCal 4.3».

The valves of mussel shells and their fragments were photographed from both the outside and the inside; the gastropod shells were photographed from different angles to reveal the diagnostically valuable traits. When identifying the shell material, we compared it with phylogenetically close recent species from the Ingoda, Onon, Shilka, and Argun rivers located near the excavation sites. The size of the fossil shells was restored by comparing their fragments with the hinges or muscle scars on valves from live collected specimens. In total, 257 shells and shell fragments from eight archaeological sites in the Upper Amur basin were collected and investigated. The examined material is kept in the collection of the Transbaikal State University (TSU; Chita, Russia) and in the scientific collection of the Institute of Natural Resources, Ecology and Cryology of the Siberian Branch of the Russian Academy of Sciences (INREC SB RAS), Chita, Russia.

Abbreviation used: tya = thousand years ago.

## RESULTS

As a result of examination of the shell collection, eight species of bivalve molluscs, including one marine species, and two species of gastropods belonging to gill-breathing and pulmonate snails, were identified. All these species are represented in the recent fauna though not all of them live now in the Transbaikalia waterbodies. A systematic list of taxa, followed by extensive comments on each identified species, is given below.

Class **BIVALVIA**

Family **Margaritiferidae** Henderson, 1929

Genus ***Margaritifera*** Schumacher, 1816

***Margaritifera dahurica*** (Middendorff, 1850)

Family **Unionidae** Rafinesque, 1820

Subfamily **Unioninae** Rafinesque, 1820

Genus ***Nodularia*** Conrad, 1853

***Nodularia douglasiae* (**Gray in Griffith & Pidgeon, 1833)

Genus ***Lanceolaria*** Conrad, 1853

***Lanceolaria* cf. *grayii*** (Gray in Griffith & Pidgeon, 1833)

Subfamily **Unioninae** Rafinesque, 1820

Tribe **Anodontini** Rafinesque, 1820

Genus ***Cristaria*** Schumacher, 1817

***Cristaria plicata*** Leach, 1814

Genus ***Sinanodonta*** Modell, 1945

***Sinanodonta schrenkii*** (Lea, 1870)

Family **Limnocardiidae** Dall, 1908

Subfamily **Adacninae** Dall, 1908

Genus ***Monodacna*** Eihwald, 1838

***Monodacna* cf. *polymorpha*** (Logvinenko et Starobogatov, 1967)

***Monodacna* cf. *colorata*** (Eichwald, 1829)

Family **Glycymerididae** Newton, 1916

Subfamily **Glycymerididinae** Dall, 1908

Genus ***Glycymeris*** (Sowerby III, 1889)

***Glycymeris* cf. *yessoensis*** (Sowerby III, 1889) – transported to area

Class **Gastropoda**

Family **Viviparidae** Gray, 1847

Genus ***Amuropaludina*** Moskvicheva, 1979

***Amuropaludina praerosa*** (Gestergildt, 1859)

Family **Planorbidae** Rafinesque, 1815

Genus ***Planorbis*** Geoffroy, 1767

***Planorbis planorbis*** (Linnaeus, 1758)

### Margaritifera dahurica

The earliest shell remains of Pearlmussels (Margaritiferidae) in Transbaikalia, dated to the late Jurassic, were found in the coal seam of the Apsat deposit. These fossils were represented by shell imprints [36]. In the Neoleistocene, the existence of pearl mussels in the central Transbaikalia was affirmed by findings made in the basin of the Upper Amur; the shells were found in the lower part of the alluvium section of the fourth terrace of the river Ingoda at the village Kaydalovo near the city of Chita [27]. The age of these finds was defined as Late-Middle-Pleistocene. The mollusks were identified as *Margaritifera* aff. *dahurica*, rather similar to the modern representatives of this species. Further data on the fossil record of Pearlmussels in the Upper Amur basin was obtained from archaeological excavations of sites and ancient settlements of Transbaikalia, where *M. dahurica* shells were the most common and numerous. In particular, fossil shells of this mussel were discovered on the territory of an archeological site situated on a high hill on the right bank of the river Ingoda, 4.5 km below Darasun settlement (Fig. 1B, 2). This site was discovered in 1975 (Fig. 2A), the cleaning of the above mentioned plot was carried out in 2019. Fragments of shells were randomly scattered over the settlement site (Fig. 2C, D). In excavation No. 3, which is 1×1 m in size, in the subsoil layer, small and large fragments of shells were relatively concentrated (Fig. 2E). Their lengths ranged from 3-5 to 9-13 cm, the total length of fossil shells, restored by comparing fragments with intact valves of the modern Pearlmussels, could reach 16–20 cm (Appendix Fig. S1). The radiocarbon age of *Margaritifera* shells from this site was 1620±60 years, and the calibrated (calendar) age was 1510±70 years.

**Fig. 2.**
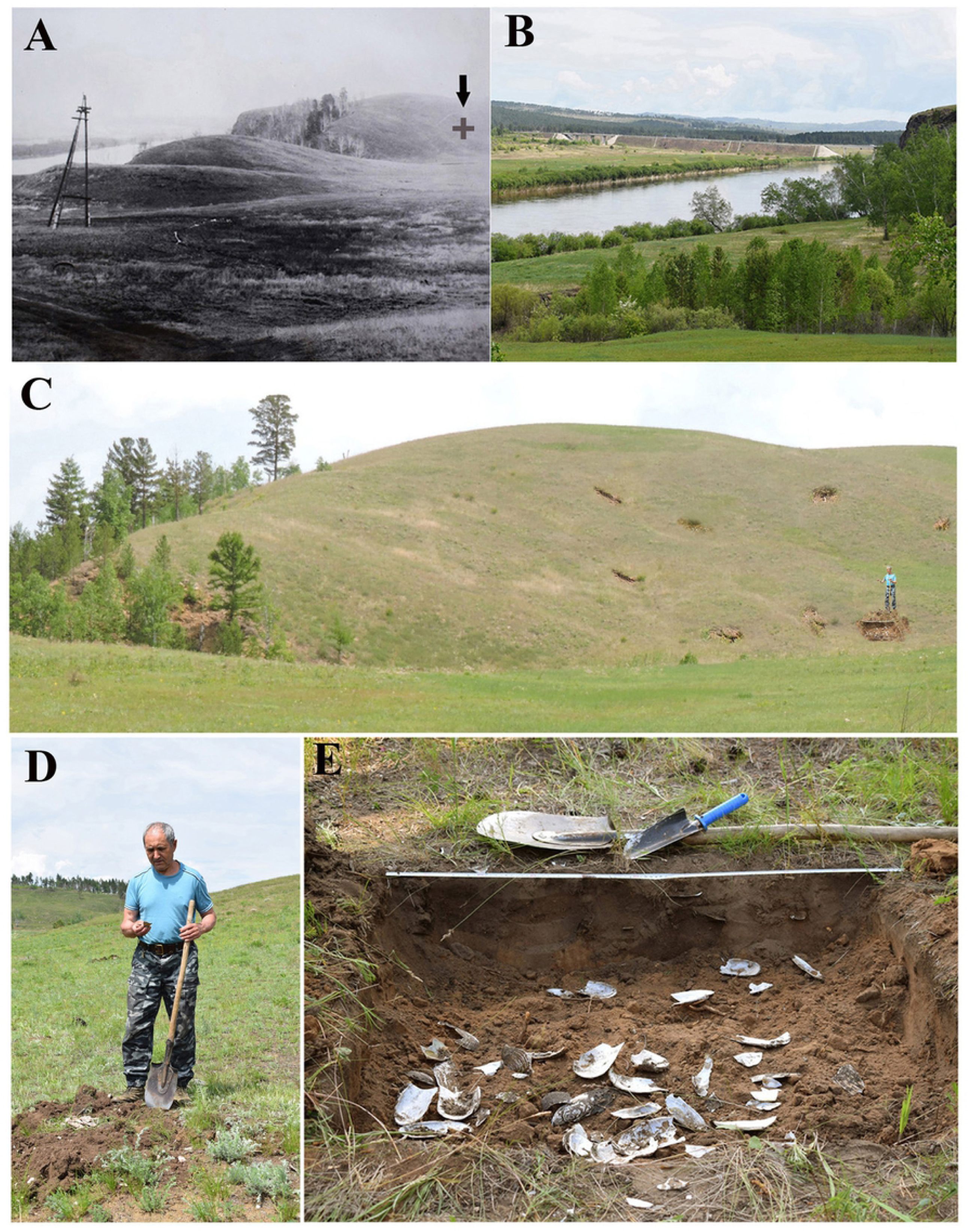
Excavations deposits on territory of the site on a hill of the right bank of the Ingoda River lower Darasun settlement. A – location of this site by data 1975 (marked by arrow **+**), B – appearance that territory in 2019; C, D – trial excavations deposits in the site in 2019; E – excavation deposits 3 with the layer of pearl mussel shells.

Pearlmussel shells were also found in excavations of ancient sites and villages in the basins of the Shilka and Argun rivers (Fig. 1C3, C4, C5, D7), whose dates made by archaeologists vary from the Neolithic to the Medieval Age (Appendix Fig. S2-S4). The radiocarbon age of their shells ranged from 2080±70 to 1690±70 years.

Shells of *Margaritifera* were most numerous in the excavations of Proezhaya 1, a large ancient settlement located on the second floodplain terrace (6–9 m high) of the river Shilka, 4.2 km below Ust’-Chernaya village of the Sretensky district (Fig. 1 C4). Inside the 200 m long and 8 to 64 m wide area protected by defensive ramparts and ditches, the remains of 70 dwellings and 20 household pits were located. Dwellings were fixed in the form of quadrangular pits ranging in size from 4×4 m to 9×12 m. Bones of wild and domestic mammals, birds, fish, fragments of ceramics and shells of river mollusks were found in the occupation layers of pits (Fig. 3A). In the dwellings themselves, shells were rare, usually they were found in household pits, outside the dwellings. For instance, in the excavation of the rampart of the defensive fortifications (Fig. 3B), a layer of shells was found containing 147 fragments and whole shells of Pearlmussels and three *Lanceolaria* sp. (shown by an arrow), but in pit No. 32 only three fragments of small margaritiferid shells were excavated (Appendix. Fig. S2).

**Fig. 3.**
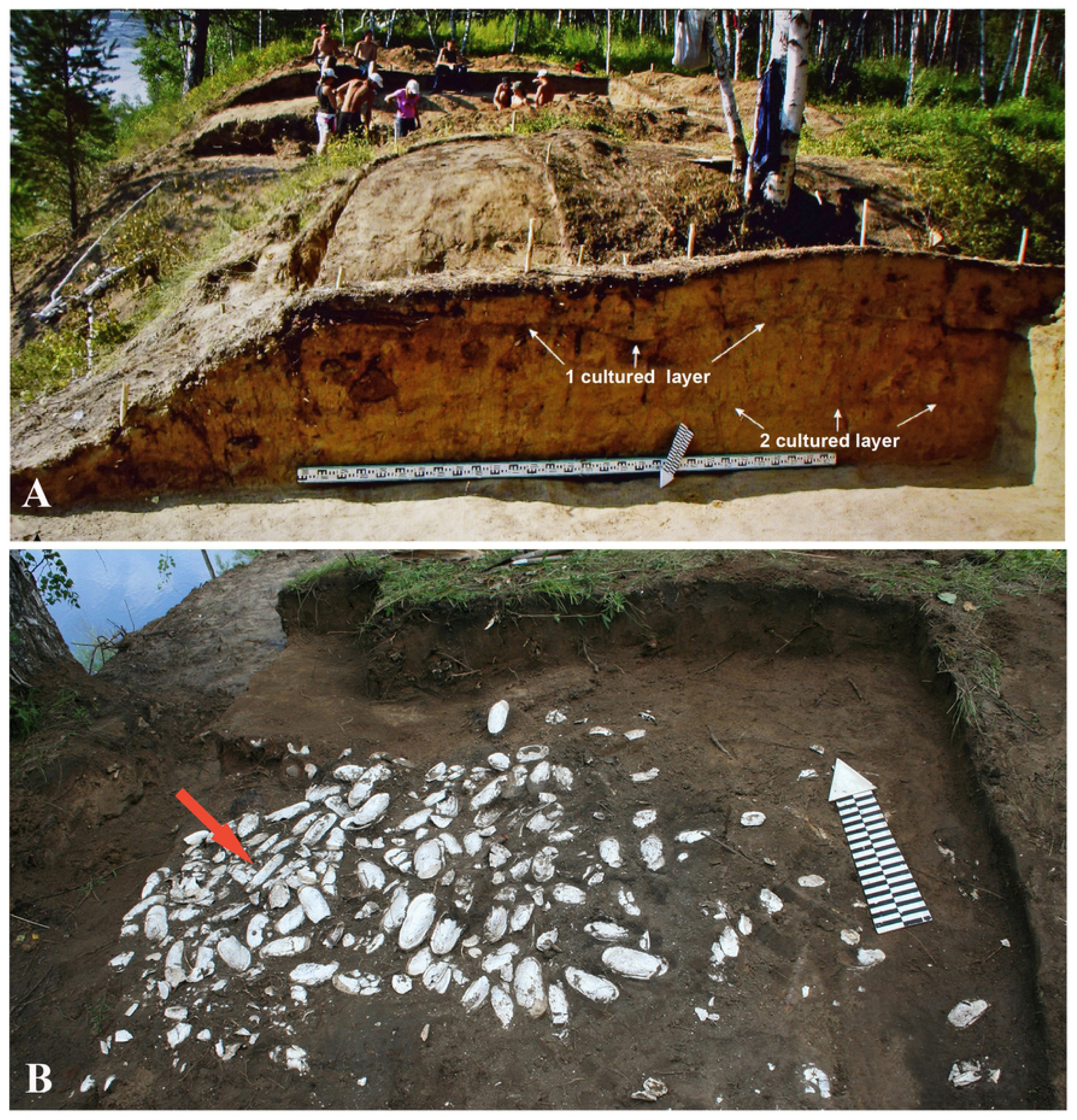
Archeological excavations in the Proezhaya 1 fortress on the bank of the Shilka River. A – stratigraphic section of the dugout wall 47, in the background – excavation deposits of the dwelling 32, B – excavation of the outside slope bank of defensive fortification with clean layer of pearl mussel shells (*Lanceolaria* shells indicated by red arrow).

### Lanceolaria grayii

A rather large, narrowly elongated *Lanceolaria* shell was found in the ritual burial of a dog inside the dwelling 28 of the ancient settlement Proezhaya 1. A juvenile dog skeleton was discovered during the excavation of the dugout floor. Shells of mollusks laid on both sides of its skull, and in front of it was an iron arrowhead (Fig. 4A), which apparently represented ritual objects placed in the burial of a guard or hunting dog. It was assumed that the dead dogs were left in the dwellings intentionally to “guard” their inhabitants from intruders. Such burials of dogs were found in several dwellings of this settlement [34]. The radiocarbon age of the shell from the dog burial is 1550±80 years.

**Fig. 4.**
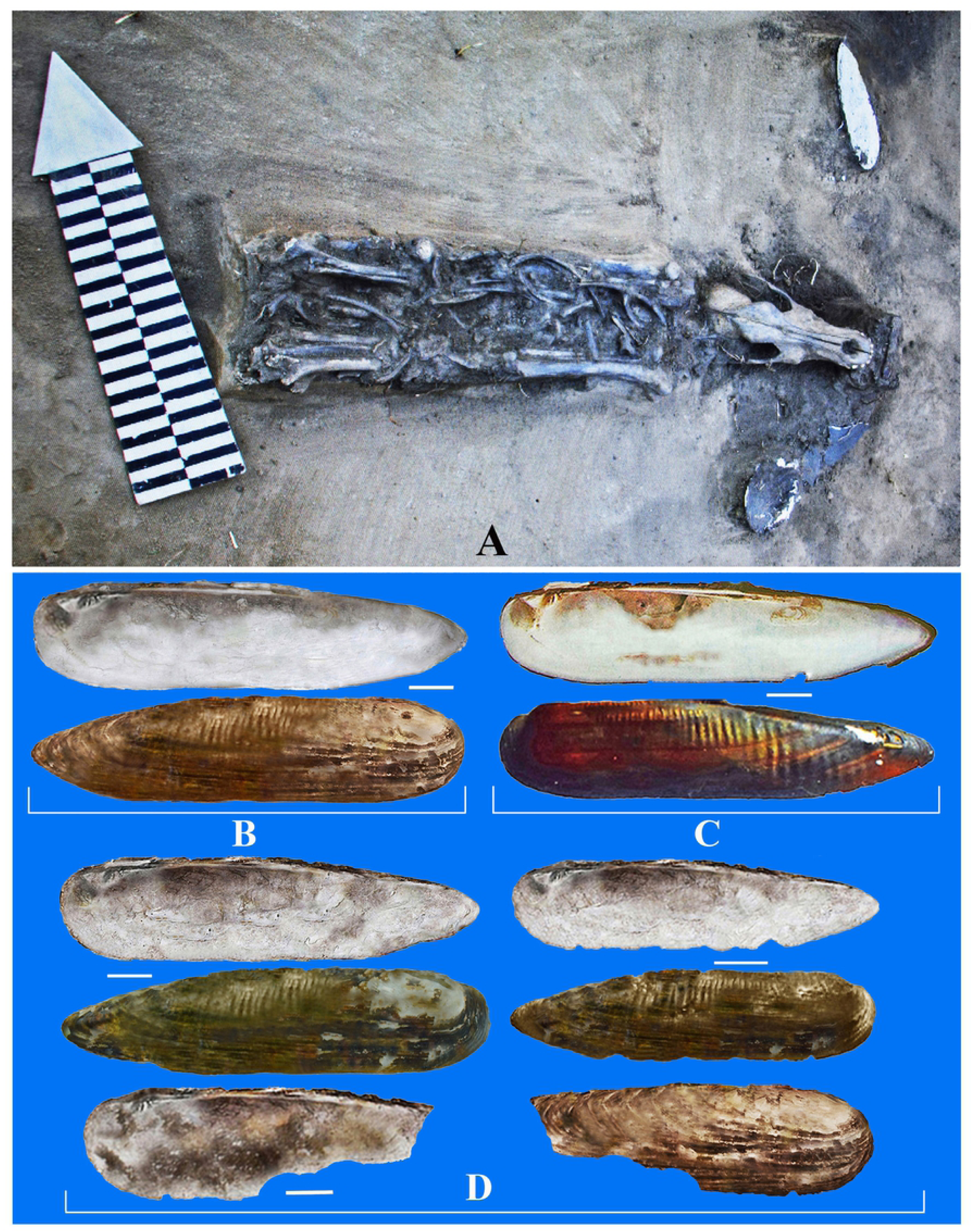
Ritual burial of the dog in the dugout 28 in the fortress Proezhaya I. A – the dog skeleton, the left of his scull is situated shell valve of *Lanceolaria* sp., to the right of it – valve fragment of *Margaritifera*. B – *Lanceolaria* cf. *grayii* from the burial of the dog, C – recent *Lanceolaria chankensis* from Khanka Lake. D – shell valves of *L*. cf. *grayii* from the bank of defensive fortifying excavation deposits. Scale bar 1 cm.

The valve of the *Lanceolaria* shell from the dog burial turned out to be identical in shape and hinge morphology with the modern *Lanceolaria grayii* from Lake Khanka (Fig. 4B-C), as well as shells from the fortification ramparts (Fig. 4D). These archaeological finds from the Shilka river basin were identified as *L. grayii*. According to molecular genetic studies, living mussels of the genus *Lanceolaria*, found in the Lake Khanka, Ussuri River basin, and Lower Amur, belong to this species [37], [38].

### Nodularia douglasiae

A fragment and an intact shell valve of *Nodularia* 4.7 and 5.9 cm long were found during the excavations of the Neolithic settlement 2 of Bol’shaya Kanga, in the Argun River basin (Fig. 1D7). The radiocarbon age of shells from this excavation was 2080±70 years.

Based on the complete morphological likeness of the shape, pseudocardinal and lateral teeth of shells from archaeological finds (Fig. 5A-D) and modern *Nodularia douglasiae* from the Argun and Ingoda rivers (Fig. 5E-I), these archaeological remains were identified as belonging to *Nodularia douglasiae*. This mussel species is widespread throughout the Amur basin, from the Magadan Region in the north to Sakhalin Island and the Primorye Territory of Russia in the south; it occurs also in Southeast Asia [16], [38].

**Fig. 5.**
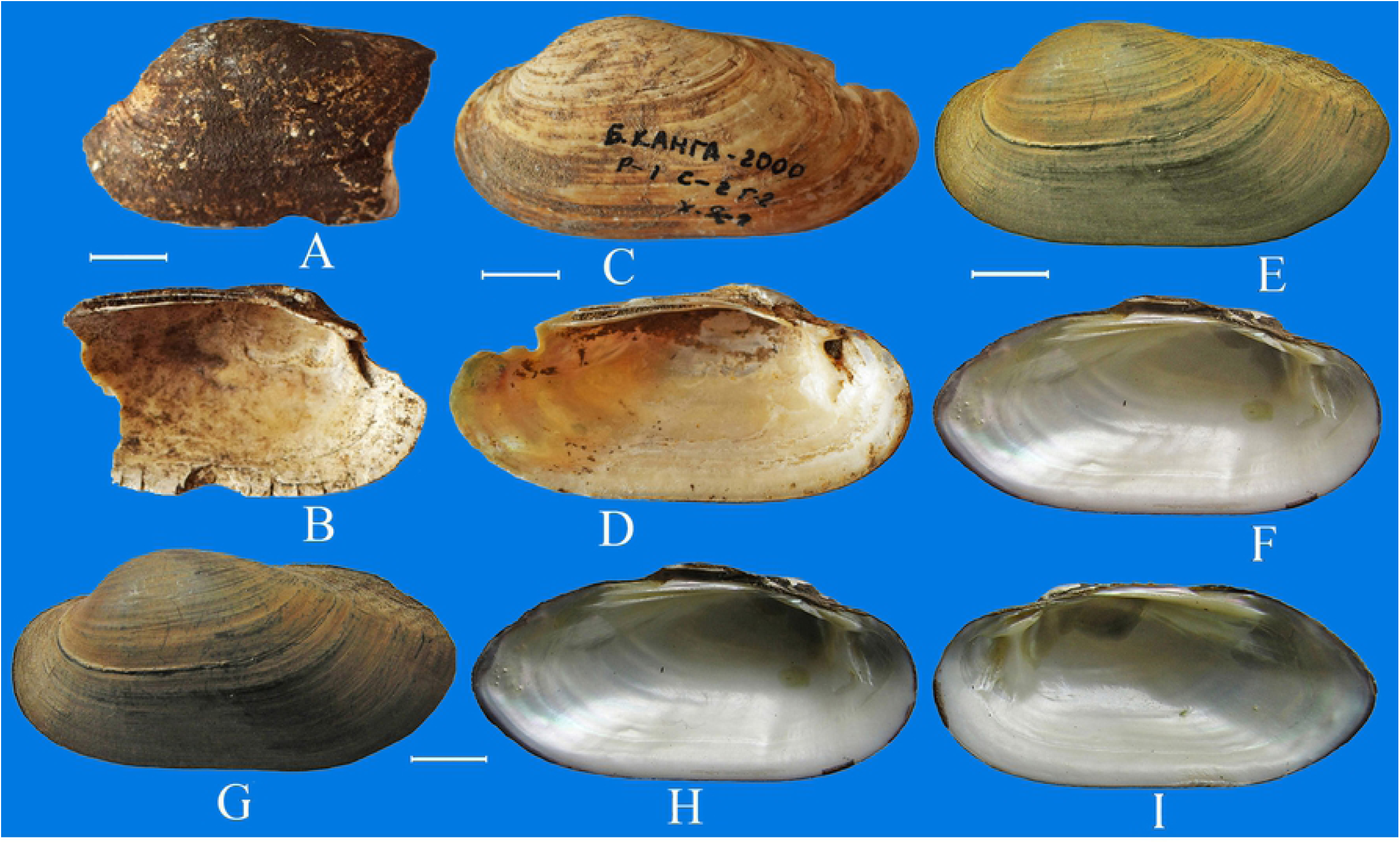
Fossil and recent shells of *Nodularia douglasiae*. A-D – shells (view on the outside and on the inside) from excavation of the Bol’shaya Kanga ancient settlement (late Stone Age), E, F – recent *N. douglasiae* from the Argun River and G-I – from the Ingoda River. Scale bar 1 cm.

### Cristaria plicata

Scattered fragments of the anodontine *Cristaria* shells were collected from excavations in the Neolithic village of Bol’shaya Kanga, Argun River basin (Fig. 1D7), and archeological site on the right bank of the Polovinka River near Luzhanky settlement (Fig. 1C3). Both archaeological finds date from the Neolithic (7-3 thousand years ago). The radiocarbon age of the shells from these excavations was 2080 ± 70 and 1690 ± 70 years.

Reconstruction of shell valves was performed by combining fossil fragments (Fig. 6A-B) with valves of recent *Cristaria plicata* from the Shilka River (Fig. 6E-F) and the Onon River basin (Fig. 6G-H). The lengths of fossil anodontine shells could be 21–29 cm in accordance with the sizes of valve fragments with lateral tooth and the size of the same portion of the shell of modern *C. plicata* (Fig. 6E-F, I-J). Now this species is widespread in the tributaries of the Amur River, in the Ussuri River basin and Lake Khanka (Russia). It occurs also in the Lake Buir-Nur (Mongolia) and countries of Southeast Asia [14].

**Fig. 6.**
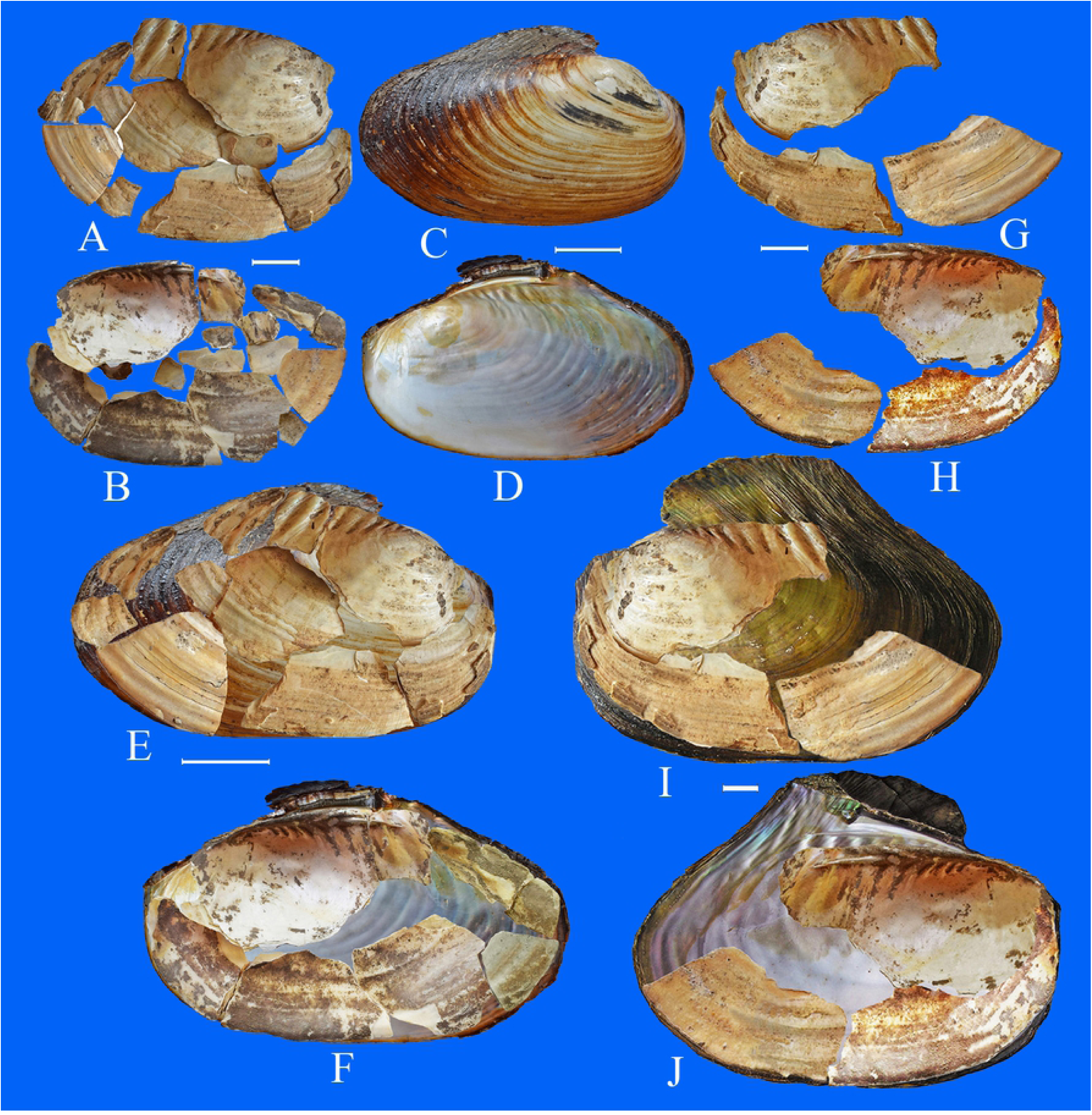
Archeological specimens and recent shells of *Cristaria*. A, B – fragments of shell valves (view on the outside and on the inside) from archeological excavation in the Bol’shaya Kanga, G-H – shell fragments of *Cristaria* from excavation in Luzhanky ancient settlement; C, D – recent *C. plicata* (view on the outside and on the inside) from the Shilka River; E-F – shell fragments from excavation in the Bol’shaya Kanga, superposed with resent shell of *C. plicata* from the Shilka River; I-J – shell fragments of *Cristaria* sp., superposed with recent shell of *C. plicata* from the Onon River Basin. Scale bar 2 cm.

### Sinanodonta schrenkii

Numerous intact shells and small shell fragments of the anodontine bivalve of the genus *Sinanodonta* together with abundant shells of Pearlmussels were found in excavations of a site situate opposite Luzhanky settlement (Fig. 1C**3**), an ancient settlement near Zheltuga river outlet (Fig. 1C**5**) and the Bol’shaya Kanga settlement (Fig. 1D**7**). The radiocarbon age of these shells was defined as 1690±70, 1770±90 and 2080±70 years, respectively. Size of fragments and intact shell valves varied from 8 cm to 15 cm (Fig. 7A-C).

**Fig. 7.**
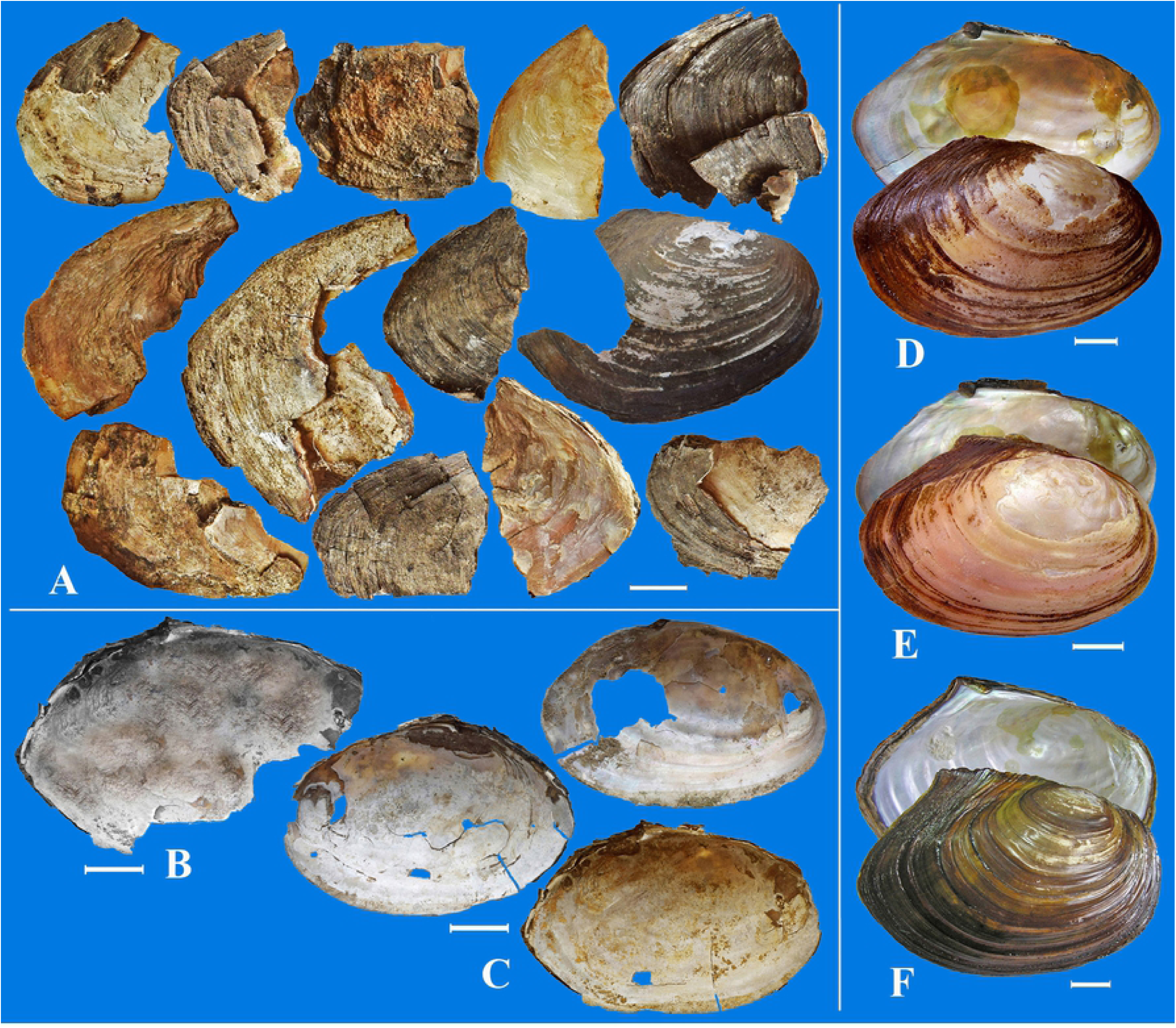
Archeological specimens and recent shells of *Sinanodonta*. A – shell fragments of *Sinanodonta* from excavation in the Luzhanky settlement, B – from the Zheltuga settlement, C – from the Bol’shaya Kanga settlement. Recent shells of *S. schrenkii*: D – from the Shilka River, E – from the Nercha River and F – from the Ingoda River. Scale bar 2 cm.

Since the excavated shells and shell fragments were morphologically similar to recent shells of *Sinanodonta schrenkii* from the Amur Basin, we identified them as belonged to this bivalve. According to molecular genetics analyses, five nominal species of the genus *Sinanodonta* inhabit the Amur River Basin and the Primorye Territory of Russia are conspecific and represent the intraspecific forms of a single valid species, *S. schrenkii* [37], [38], which is widespread in the Amur River Basin and the Primorye Territory of Russia.

### *Monodacna* cf. *polymorpha*

#### *Monodacna* cf. *colorata*

Fossil shells of two cardiid bivalves of the genus *Monodacna* were found in the floodplain of the Ingoda River, on the territory of Antipikha settlement, a suburb of Chita City (Fig. 1B**1**). The width of the floodplain in this place reaches 1.5 km; the site where the shells were found is situated in 500 m from the current riverbed (Fig. 8A). One shell was found in 2009 during making a well; three valves of *M*. cf. *polymorpha* and one of *M*. cf. *colorata* (valve length 16–18 mm) were collected in 2019 at the same place and from a soil drillhole (diameter 16 cm, depth 6-8 m). The shells, morphologically similar with recent *M. polymorpha* and *M. colorata*, are rather hard, convex, ovate-triangular, with narrow smoothed umbo, large triangular cardinal tooth, and shallow muscular scars and mantle sinus. Shell surface of *M*. cf. *polymorpha* is covered by narrow and frequent smoothed ribs, the width of which exceeds the width of the interrib intervals (Fig. 8B). The ribs on shells of *M*. cf. *colorata* are wider, with acute not smoothed ribs (Fig. 8C). The absolute age of the fossil bivalves cannot be determined by radiocarbon dating but it must surely exceed 50 tya (for details see *Planorbis planorbis* section and Discussion). The genus *Monodacna* is a part of the Ponto-Caspian malacofauna; its recent representatives live only in the basins of the Black, Caspian, and Aral Seas [39], [40].

**Fig. 8.**
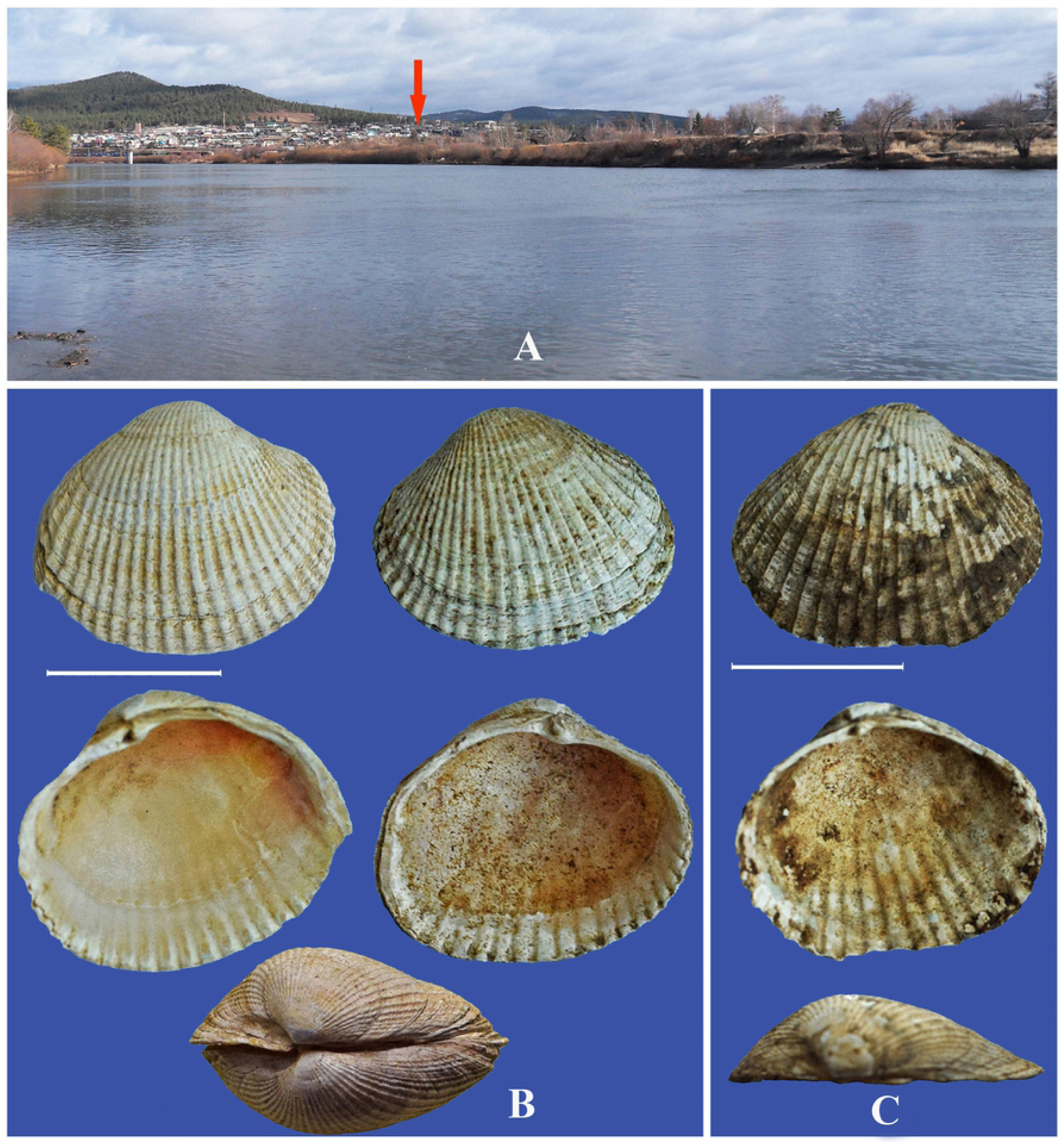
Locations of *Momodacna* species. A – Ingoda River near Antipikha settlement, suburb of Chita city. Fossil shells (view on the outside, on the inside and the above) from drill hole in the Ingoda River flood land: B – *Monodacna* cf. *polymorpha*, C – *Monodacna* cf. *colorata*. Scale bar 1 cm.

### *Glycymeris* cf. *yessoensis*

*Glycymeris yessoensis* belongs to marine bivalve mussels of the family Glycymerididae. A shell of *Glycymeris* cf. *yessoensis* was found in 1991 in the burial ground on a terrace-like plateau near Duroyskiye lakes in the Argun River Basin (Fig. 1D**8**). In total, 12 archaeological sites were discovered in this area. Among them – the Paleolithic settlement of Duroy, belonging to the 1^st^ half of the Sartan glaciation (25–13 thousand years ago), where bones of a woolly rhinoceros, bison, mammoth, reindeer and stone tools were found. In the burial ground dated to the late Neolithic – Early Bronze Age, under the oval stone mounds of a diameter of 8-9 m, in grave pits 2.5-3 m deep, tribes living in the 3^rd^ century BC–2^nd^ century AD buried their deads [41]. In the burial ground with funerary equipment, including fragments of clay vessels, stone products, animal bones, and shells of river mollusks. Shell of *Glycymeris* cf. *yessoensis* 42 mm long, with a hole in the umbonal area made for the sake of using it as a pendant, was discovered. The radiocarbon age of the shell was 5540 ± 160 years, the calibrated age was 4820 ± 130 years.

According to the overall shell morphology, the shape of muscle scars and the mantle sinus, the shells from the Duroy burial ground is similar to recent *G. yessoensis* from the Amursky Bay (Sea of Japan/East Sea) (Fig. 9A, B). The shell of *G. yessoensis* from the excavation of the medieval settlement Nikolaevskoe I in Primorye [44] also resembles the shell from the Duroy burial site, but differs by the symmetrical structure of the valves and the location of the mantle sinus slightly above the ventral edge (Fig. 9C). In samples from the Duroy burial ground and the Amursky Bay, the posterior margin of the shell is slightly longer than the anterior one; the mantle sinus is located very close to the ventral margin of the shell. *Glycymeris yessoensis* is known as a common species of the South Kurile shallow waters, found in the Amursky Bay (Sea of Japan/East Sea) [42], [43].

**Fig. 9.**
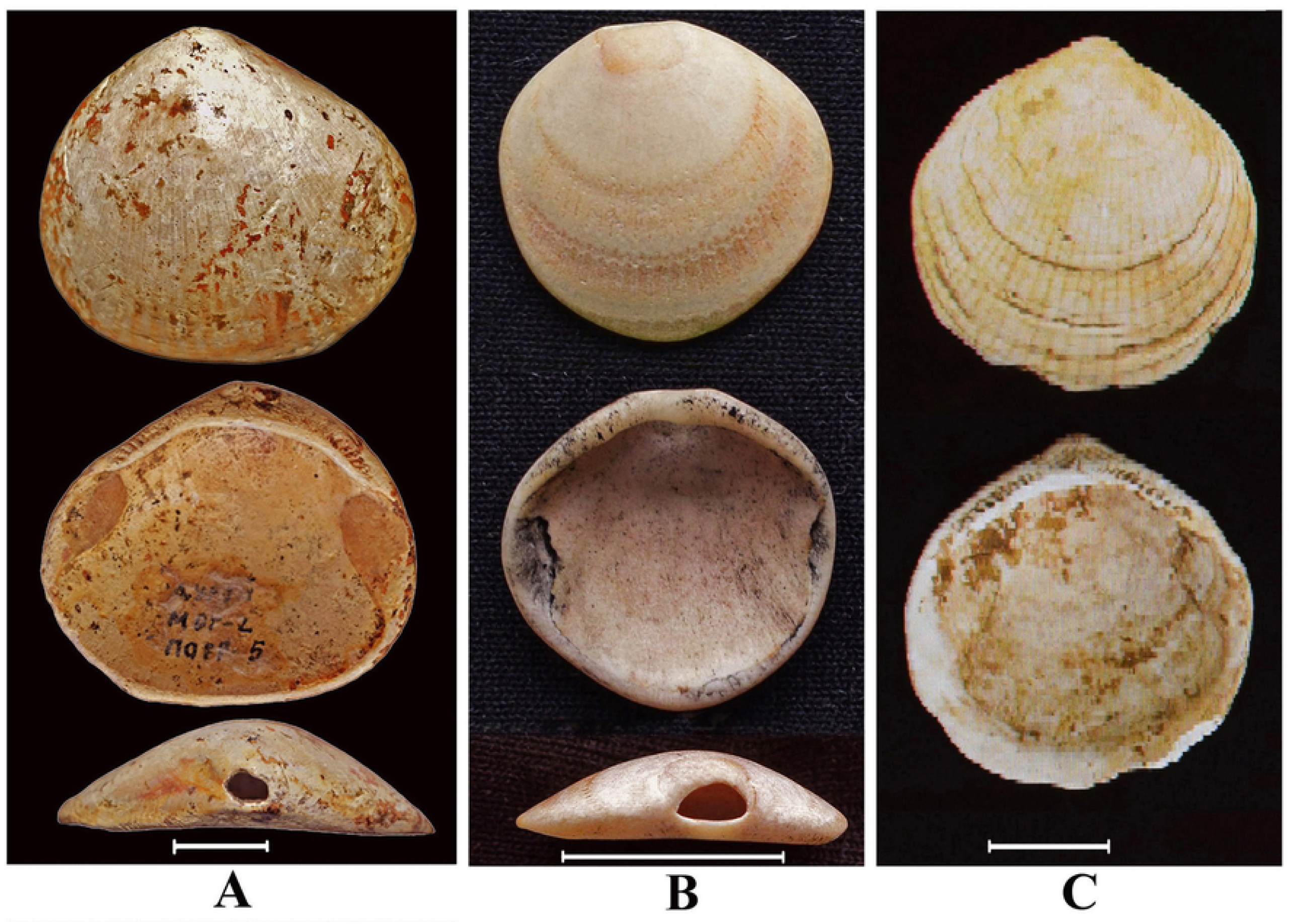
Archaeological specimens and recent shells of *Glycymeris*. View on the outside, on the inside and the above: **A** – *G*. cf. *yessoensis* from the Duroy burial ground in Transbaikalye, **B** – recent *G. yessoensis* from the Amursky Bay (Sea of Japan/East Sea), the hole in the umbonal area of valve (view on the above) is a result of activity of a sponge drilling; **C** – *G. yessoensis* from excavation of the medieval settlement Nikolaeskoe I in the Primorye, Russian Far East [44]. Scale bar 1 cm.

### *Amuropaludina* cf. *praerosa*

A fossil gastropod shell morphologically similar to shells of the recent *A. praerosa* from the Amur river was discovered in the burial ground of Zorgol in the burial 6/4, dated I century. BC – I century AD. The absolute age of this shell by radiocarbon dating is 2160 ± 90 years. The Zorgol burials in the Priargunsky area (Fig. 1D6) are represented by oval stone depositions, with grave pits of a rectangular shape 2–3 m deep. The burial items included clay and birch bark dishes, bone and iron arrowheads and spears, horn covers of bows, and various adornments [45]. Bones of sacrificed animals and shells of aquatic mollusks were found in the fill of burial pits and inside burials. The thick-walled high-turreted shell found in the burial, 31 mm high and 22 mm wide, had two holes in the upper whorl, which appear to be made for the sake of using it as a pendant (Fig. 10A-D).

**Fig. 10.**
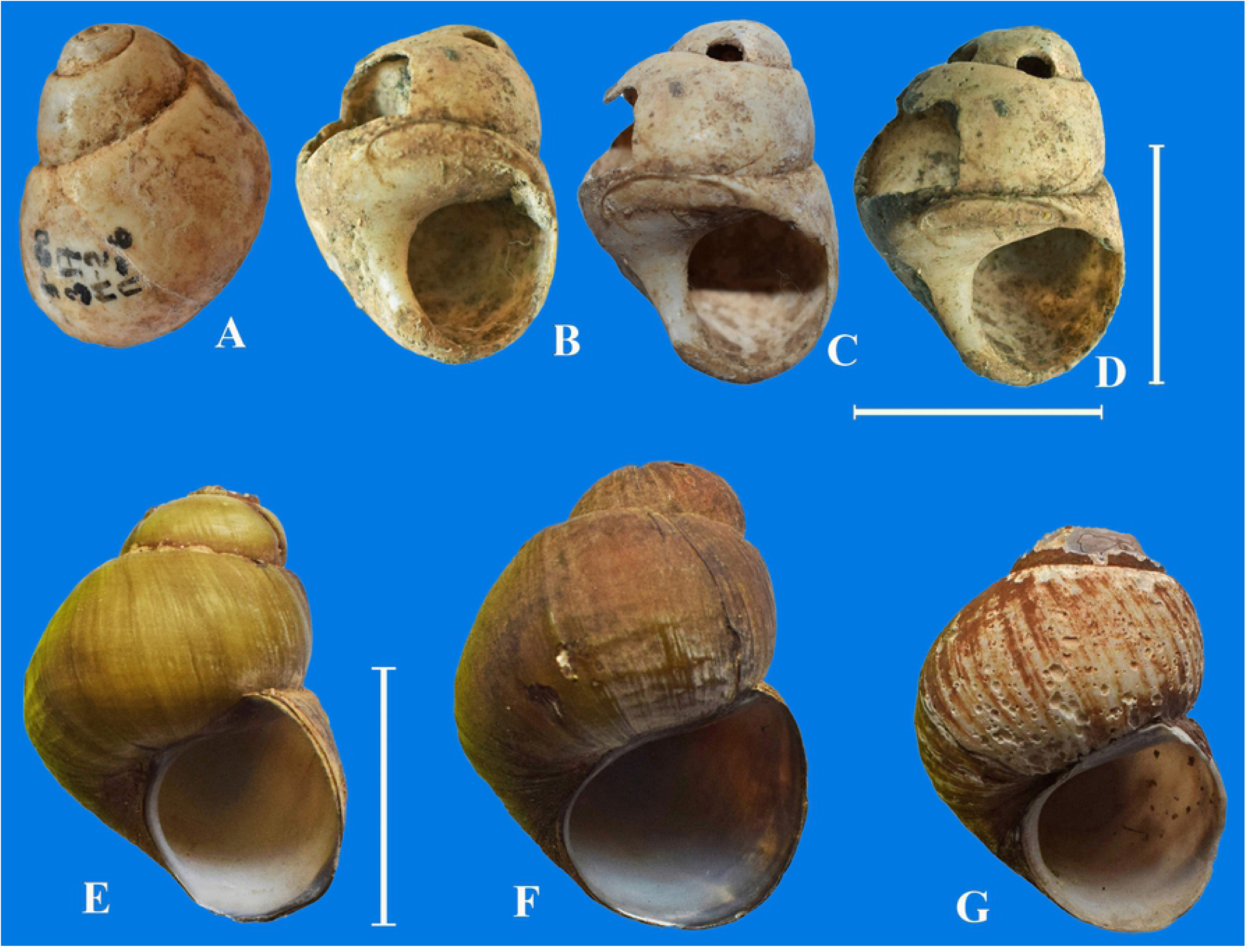
Shells of *Amuropaludina praerosa*. A-D – view of fossil shell *A*. cf. *praerosa* in different perspective from Zorgol burial ground, E-G – recent shells of *A. praerosa* from the Middle Amur River. Scale bar 2 cm.

Living *A. praerosa* of similar size were collected in 2006 from the Middle Amur River near Poyarkovo settlement (Fig. 10E-G). Currently, the distribution of this species is limited to the middle and lower parts of the Amur basin [39]. It does not occur in the Transbaikalia area [46].

### Planorbis planorbis

The fossil shell of the gastropod *Planorbis planorbis* from the alluvial deposits of the Ingoda River was extracted from a borehole located in vicinities of Chita City (Fig. 1A1). This find was accompanied by shells of the cardiid genus *Monodacna* (see above). This shell bears all characteristics of Recent representatives of this species: it is rather small, discoidal, covered by fine axial striations. Shell whorls are slightly convex and increase relatively quickly and evenly, the last whorl is approximately twice the width of the penultimate one (Fig. 11). A pronounced keel is visible along the basal surface of the last whorl, the aperture is evenly rounded, without a distinct angulations. Shell dimensions (in mm) are: width 9.8, height 2.2.

**Fig. 11.**
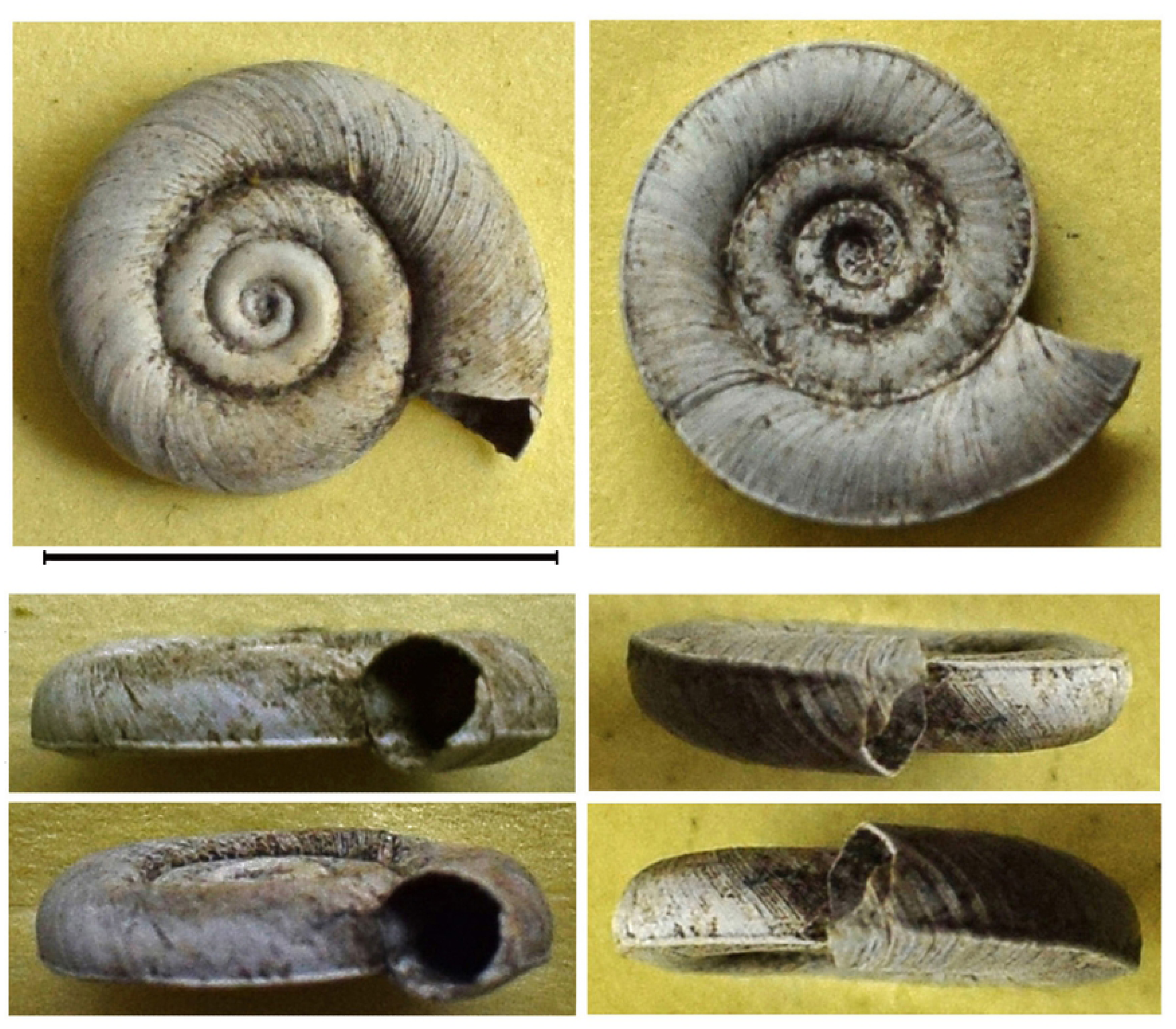
Fossil specimen of *Planorbis planorbis*. Shell from the drill hole with Pleistocene deposits in the flood land of the Ingoda River.

We were unable to determine the absolute age of shells from this locality (both *Monodacna* and *Planorbis*) because of limited resolution of the radicarbon dating method. The upper age of an object may be older than 50 tya because of a relative short half-life of the ^14^C isotope. Most probably, the existence of both *Monodacna* and *Planorbis* in Transbaikalia can be assigned to the Kazantsevsky interglacial of the Neopleistocene, in the time interval from 130–70 tya (see Fig. 12. Discussion).

**Fig. 12.**
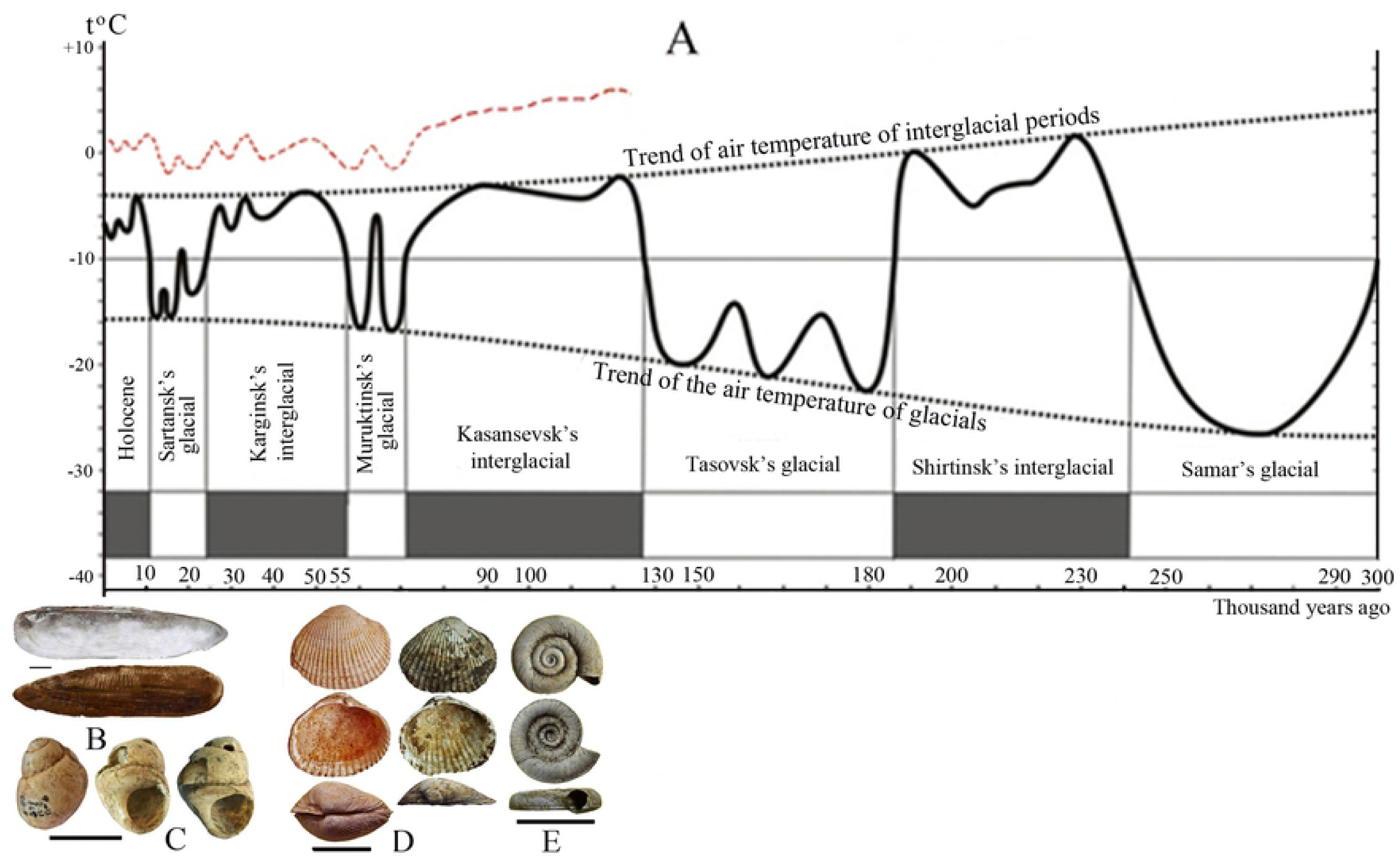
A – Change of the average annual temperature of ground air in epochs of Neopleistocene and Holocene according to palynological analyses of the core in Charskaya depression in the northern Transbaikalye [57], [28]. Red dashed line – a probable temperature trend in the Upper Amur Basin in the south-eastern Transbaikalye. Possible time of existence of thermophilic species in the Upper Amur Basin: **B** – *Lanceolaria* cf. *grayii* и **C** – *Amuropaludina* cf. *praerosa*, **D** – *Monodacna* cf. *polymorpha* and *Monodacna* cf. *colorata*, **E** – *Planorbis planorbis*. Scale bar for mussels = 1 cm.

Fossil shells of *Planorbis planorbis* are known from the late glacial lacustrine sediments of Chany Lake, south part of the West Siberian Plain [47]. This species in the Pleistocene sediments of the Kirenga River Basin in the Eastern Siberia also was found [48]. These sediments were represented by gravel pebbles with a lenses of green sands in the Kirenga River paleobed. According to Popova [48], the age of *P. planorbis* in the Baikal area should not be older than the Lower – the early Middle Pleistocene, as indicated by spore-pollen complexes from this location, represented by pollen of spruce (*Picea excelsa*), cedar (*Pinus sibirica*), pine (*Pinus silvestris*), fir (*Abies* sp.), larch (*Larix* sp.), linden (*Tilia* sp.), oak (*Quercus* sp.), elm (*Elmas* sp.) and hemlock (*Tsuga* sp.), according to data of Kultchitski et al. [49]. The recent range of *P. planorbis* covers much of Europe and Western Siberia eastward to Altai Mts. [39], [40], [50]. It is absent from the Baikal Lake basin and adjacent areas [51], though old records of this species (under the name *Planorbis marginatus*) from the shallow bays of the Baikal Lake are known [52], [53]. During faunistic surveys of freshwater mollusca of the south part of Eastern Siberia, including Lake Baikal, this species has been not found (M. Vinarski, pers. observ.). It is reasonable to assume that *P. planorbis* is extirpated now in the studied area.

### The Neopleistocene-Holocene molluscs as indicators of climate change

To assess the value of the studied fossil molluscs as possible indicators of the paleoclimatic conditions, we compared the absolute age of identified species of gastropods and bivalves and the climatic phases (climatochrones) of the regional geochronological scale. In the Holocene era (Mesolithic-Iron Age period), between 10.0–2.0 tya, four climatic periods were distinguished from the Boreal to the Subatlantic (Table). In the Neopleistocene (Paleolithic-Early Mesolithic), the Sartansky cryochron (25.0–10.8 tya) included 4 cryostages and 3 thermal stages. The early Paleolithic Age of Transbaikalia coincided with the Karginsky thermochron dated 50–25 tya. According to archaeological data and radiocarbon dating, the period of presence of the studied mollusc species in the Upper Amur basin does not coincide with the stages of the regional climatochronological scale.

Obviously the paleogeographic situation in the southeast of Transbaikalia (50-52° N, 114-120° E) with a low-mountainous relief was significantly different from that in its northern high-mountainous regions (56.5° N), where powerful glaciation occurred in the Neopleistocene [57]. According to data of Dr. F. Enikeev (Institute of Natural Resources, Ecology and Cryology SB RAS, personal communication), there was no a continental glacier in the southeast of the region, on the territory of the Upper Amur basin, but there were huge ice-dammed reservoirs interconnected with the vast glacial basins of Eurasia. The climate changes in this area were more smooth than in the north of the region.

According to paleontological data, the Pearlmussels (Margaritiferidae) in the Pleistocene of central Transbaikalia existed during cold climatic phases [27]. The archaeological findings show that *Margaritifera dahurica* lived in the southeast of Transbaikalia, in the Upper Amur basin from the Sartansky cryochron to the Holocene thermochron. This characterizes *M. dahurica* as an eurythermic species. *Lanceolaria grayii* and *Amuropaludina* cf. *praerosa*, according to radiocarbon dating, occurred in the Upper Amur basin during the Holocene thermochron, and disappeared here during the late Holocene cooling, when the average annual air temperatures dropped below the optimum (+1.0–1.8° C), at which these species currently live in the Far East of Russia (46-52° N, 130-140° E). Both *Lanceolaria grayii* and *Amuropaludina praerosa* can be characterized as stenothermal molluscs. The *Monodacna* species are now confined to the Ponto-Caspian area and occur in the basins of the Black, Caspian, and Aral seas. Their current range assumes the relative thermophyly of these cardiids. *Planorbis planorbis*, with its wide distribution, is, probably, the most eurythermic species of all discussed in this paper. Its collection from northern latitudes of Europe and Western Siberia are known [54], [55], [56] and, thus, the performance of this snail as a climate change indicator is very low. We can assume that *Monodacna* and *Planorbis* lived in the Upper Amur basin, probably during the Karginsky and Kazantsevsky interglacials of the late Pleistocene and became extinct there during the Murukta or Sartan glaciation. The fact of the existence of these species in Eastern Siberia in the Neopleistocene suggests their wider ranges with peripheral and isolated (?) populations in the Transbaikalia area. The rest of species identified by us in archaeological samples (*Nodularia douglasiae, Cristaria plicata, Sinanodonta schrenkii*) could survive in this region up to modern time, we consider them eurythermal and eurybiontic.

The amplitude of climate change in Transbaikalia during the Neopleistocene and Holocene is evidenced by palynological studies that record cycles of glaciation and deglaciation over a long period of time (up to 300 tya) [57], [28]. According to palynological analyzes of 1.2 km of core in the Chara depression in the north of Transbaikalia (56.5° N, 116° E), the average annual air temperature in the Neopleistocene varied from +1 to –4° C, in the Holocene from –4 to –8° C, and during glacial stages from –16 to –22° C. (Fig. 12). The age of the palynological complexes, confirmed by isotopic dating, shows a rather detailed picture of the dynamics of temperature indicators in the north of Transbaikalia, and the change in the species composition and the timespan of the fossil malacofauna in the Upper Amur basin reflect the dynamics of climate change in the southeast of Transbaikalia.

The significant heterogeneity of landscape and climatic conditions and the altitudinal position of the regions in Transbaikalia caused the mosaic formation of malacofauna both in the past and in the present, with a limited distribution of stenoecous species in the Upper Amur basin, which experienced less sharp climate fluctuations than the north of the region. When reconstructing the living conditions of fossil molluscs, the actualistic method of analogy based on knowledge of ecology and life habits of recent closely related species can be recommended [59], [7].

The species *Lanceolaria grayii* and *Amuropaludina praerosa*, which are now extirpated in Transbaikalia, inhabited the Upper Amur basin 1550±80 – 2180±90 years ago, under milder climatic conditions similar to the modern climate of the Russian Far East. Their range covered the territory from the Upper to the Lower Amur River basin, Ussuri River basin and Lake Khanka. *Monodacna* and *Planorbis* lived here in the Neopleistocene, probably from 130 to 70 tya, in a warmed shallow lake or in a flooplain waterbody of the Ingoda River, in a neutral or slightly alkaline environment, on soft silty-sandy soils, summer water temperature could be around 22–24° C (by analogy with today’s environment of waterbodies of the Ingoda River floodplain). Although most representatives of the genus *Monodacna* are brackish water species, the modern invasion of *Monodacna colorata* in the lower and middle reaches of the Volga [39] shows that this species is not stenohaline and can exist under strictly freshwater conditions.

These molluscs, as indicator species, provide some information on climate change in the southeast of Transbaikalia during the Holocene and the Neopleistocene.

## DISCUSSION

Shells and shell fragments of the pearl mussel *Margaritifera dahurica* were the most numerous among the mollusc remains found during archaeological excavations in Transbaikalia. The massive aggregations of its shells in archeological sites, ancient households and hillforts (Fig. 2; Appendix Fig. S2-S4) suggest that the prehistoric humans collected these mussels mainly as a food item. The shells were also used for handicrafts and jewelry made of mother of pearl (buttons, arrowheads and spearheads, spinners, disk-shaped beads and bracelets) (Appendix Fig. S5).

The species of the genera *Lanceolaria, Nodularia, Cristaria* and *Sinanodonta* are much rarer (Fig. 4B-D, 5A-D, 6A-B, 7A-C), though the ancient people could also use them for food and as a material for crafts and decorations. The finding of pendant made of *Amuropaludina* cf. *praerosa* shell (Fig. 10A-D) in the Zorgol burial ground indicates that this species inhabited the Argun River Basin nearly 2.1-1.6 tya in a milder climatic conditions as compared to the current ones.

An interesting find was a shell of the marine bivalve species, *Glycymeris* cf. *yessoensis*, in the Duroy burial ground (Fig. 9A). It also was used as a pendant, which, perhaps, served as an offering and then a ritual object during the burial. The shell of *Glycymeris* cf. *yessoensis* from the burial place turned out to be quite similar to shells of *Glycymeris yessoensis* from the excavation of the medieval Nikolayevskoye 1 settlement in Primorye. Possibly, the ancient tribes of the Amur and Primorye areas had transport and trade connections, as a result of which the shell of *Glycymeris* (of non-local origin) could have appeared in Transbaikalia. This serves as an important complement to archaeological information about the lifestyle and culture of the ancient people of Transbaikalia.

The identification of shells of *Monodacna* and *Planorbis* (Fig. 8B-C and Fig. 11) from a borehole located in the Ingoda River floodplain is unique for Transbaikalia since these molluscs do not occur here anymore. Their current ranges are situated much to the west, in the southern part of Europe and Central Asia (*Monodacna* and *Planorbis*) and Western Siberia (*Planorbis*). It shows that their ranges were relatively recently much wider. Possibly, the isolated parts of their ranges were located in Transbaikalia. Though the disappearance of the cardiid clams from the Baikal area may be easily explained by climate cooling and the disruption of the former interbasin connections, the regional extirpation of *Planorbis* remains somewhat enigmatic. This snail is tolerate of cold climate and is able to live farther north than in Transbaikalia.

According to archaeological and radiocarbon dating, the existence of extirpated mollusc species in the southeast of Transbaikalia does not correspond with the stages of the climatic timescale developed for the whole region (Table). It possibly indicates the different paleogeographic situation and the magnitude of climate change in the Pleistocene and Holocene in particular subregions of Transbaikalia.

During the periods of the Neopleistocene and Holocene climatic optima, in the Upper Amur basin, on the territory of Transbaikalia, five regionally extirpated mollusc species lived (Fig. 12). Three of them (*Monodacna* cf. *colorata, M*. cf. *polymorpha*, and *Planorbis planorbis*) disappeared in the Neopleistocene, whereas *Lanceolaria grayii* and *Amuropaludina praerosa* were present in the region until the mid-Holocene. In our opinion, these species may be considered more or less reliable indicators of the climate change in the Transbaikalia region.

Changes in the species composition of the malacofauna of Transbaikalia associated with long- and short-term natural cycles of warming in the interglacial and cooling periods during the Neopleistocene and Holocene can be considered as a manifestation of climatogenic succession. During this succession, there was a decrease in the species diversity of mollusks at the regional level and a reduction in the peripheral areas of the ranges of thermophilic species at the global level.

## Acknowledgements

Authors would like to thank Dr. F.I. Enikeev (Institute of Natural Resources, Ecology and Cryology SB RAS) for useful discussions about climate change in Transbaikalye in glacial periods of the Neopleistocene and Holocene. The authors also would like to thank Dr. Kh.A. Arslanov (Institute of Sciences about Earth, Saint-Petersburg State University) for the radiocarbon dating of fossil molluscs shells. Special thanks are due to I.E. Mikheev (Institute of Natural Resources, Ecology and Cryology SB RAS) for organization of field works and help with sampling of recent molluscs species in the Upper Amur Basin on the Transbaikalye territory.

---

**Table.**
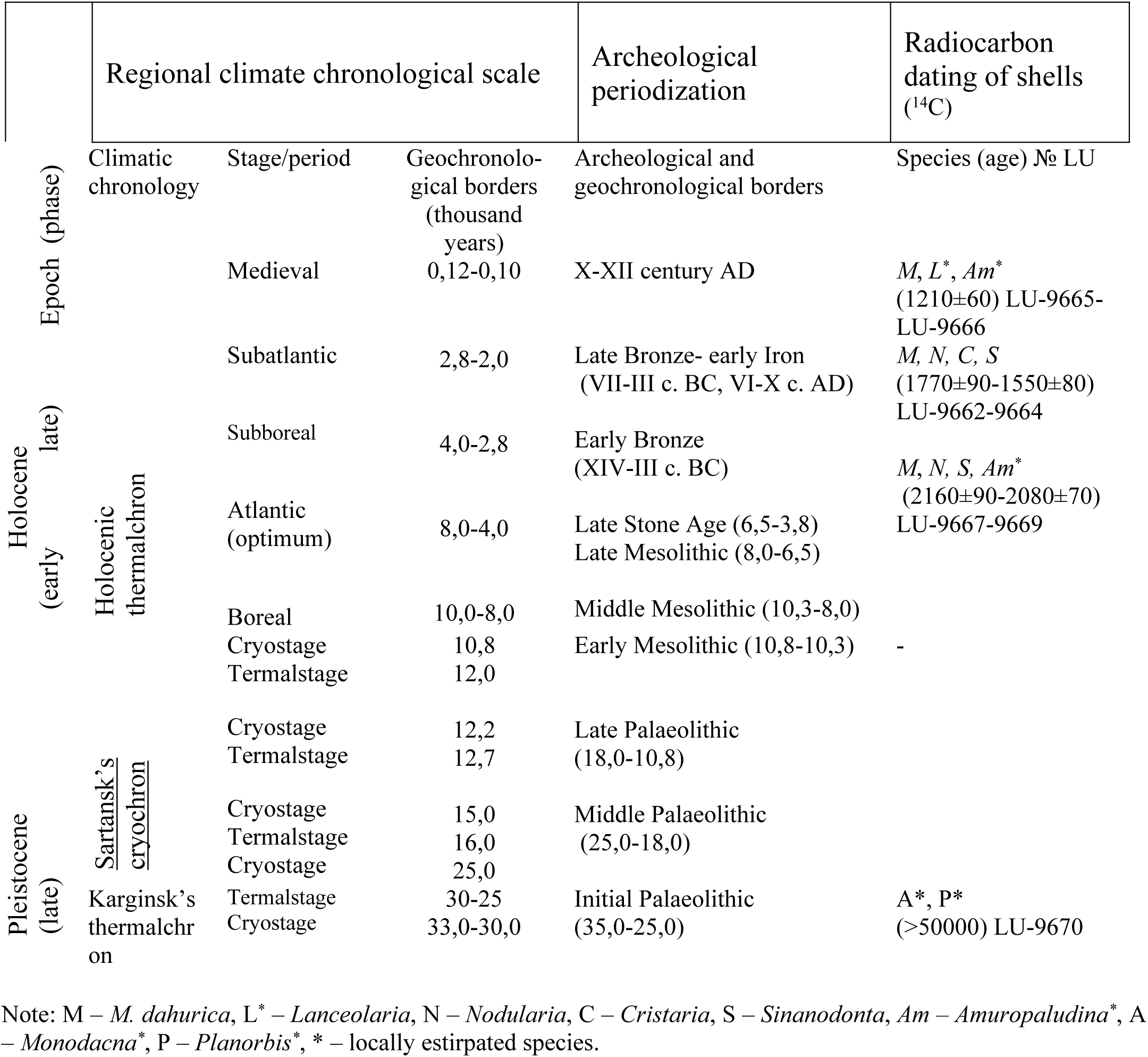
General geochronological scale for Late Pleistocene and Holocene with regional climate chronological scale [27], subdivisions of archeological periodization [60] and radiocarbon dating of the fossil molluscs (by our data)

## Figure captions for Supporting Information

**Fig. S1**. Archeological specimens of the margaritiferid mussels. Shells of *M. dahurica* from excavation of ancient site near the Darasun settlement (1620±60 years). A-C – reconstruction of shell length by overlapping of some fossil shell fragments with valves of the recent *M. dahurica* from the Ingoda River.

**Fig. S2**. Shells of *M. dahurica* from excavation of medieval Proezhaya 1 fotress (1550±80 years). A – from the bank of defensive fortifying, B – from the ancient dwelling 32 and reconstruction of fossil shells length by superposition with the recent shell valves of *M. dahurica* from the Shilka River.

**Fig. S3**. Shells of *M. dahurica* from excavation of the ancient settlement near mouth of the Zheltuga River (1770±90 years). Scale bar 2 cm.

**Fig. S4**. Fragments of valves and whole shells of *M. dahurica* from excavation of the Bol’shaya Kanga ancient settlement (2080±70 years), and reconstruction of fossil shell length by superposition with the recent shells of *M. dahurica* from the Argun River.

**Fig. S5**. Handmade goods from the shell molluscs. A – button from nacre and blanks for beads and bracelet, B – arrowheads an d spearheads, C – adornments in the form of the pendant.

## References

1. Bogan A. Global diversity of freshwater mussels (Mollusca, Bivalvia) in freshwater. Hydrobiologia. 2008; 595: 139–147.

2. Strong, EE, Gargominy O, Ponder WF, Bouchet P. Global diversity of gastropods (Gastropoda; Mollusca) in freshwater. Hydrobiologia. 2008; 595: 149–166.

3. Lydeard C, Cummings KS, editors. Freshwater mollusks of the World: A distribution atlas. Baltimore: John Hopkins University Press; 2019.

4. 108. MolluscaBase. 2020. Accessed at http://www.molluscabase.org on 2020-04-25

5. Gray J. Evolution of the freshwater ecosystem: The fossil record. Palaeogeogr Palaeocl. 1988; 62: 1–214.

6. Benton M, ed. The Fossil Record 2. London etc.: Chapman & Hall; 1993.

7. Tolstikova NB. About possibility usage of molluscs for reconstruction of paleolimnological conditions in ancient lakes of arid and damp climate. In: Martinson GG, editor. Paleolimnology of lakes in arid and humid zones. Leningrad: Nauka; 1985. pp. 62–85. (In Russian).

8. Tolstikova NB. Climatic influence of palaeolimnological and malacofauna changes in Cenozoic of Zaisan depression. In: Kvasov DD, Starobogatov YI, editors. Climate and fauna of Cenozoic. Leningrad: Zoological Institute of the Soviet Academy of Sciences; 1985. pp. 66–73. (In Russian).

9. De Francesco CG, Zárate MA, Miquel SE. Late Pleistocene mollusc assemblages and inferred paleoenvironments from the Andean piedmont of Mendoza, Argentina. Palaeogeogr Palaeocl. 2007; 251(3-4): 461–469.

10. Pisano MF, Fucks E. Quaternary mollusc assemblages from the lower basin of Salado River, Buenos Aires Province: Their use as paleoenvironmental indicators. Quatern Int. 2015; 391: 1–12.

11. Georgopoulou E, Neubauer TA, Strona G, Kroh A, Mandic O, Harzhauser M. Beginning of a new age: How did freshwater gastropods respond to the Quaternary climate change in Europe? Quaternary Sci Rev. 2016; 149: 269–278.

12. Bolotov IN, Bespalaya YV, Vikhrev IV, Aksenova OV, Apsholm PE, Gofarov MY, Klishko OK, Kolosova YS, Kondakov AV, Lyubas AA, Paltser IS, Konopleva ES, Tumpeesuwan S, Bolotov NI, Voroshilova IS. Taxonomy and Distribution of Freshwater Pearl Mussels (Unionoida: Margaritiferidae) of the Russian Far East. PLoS ONE. 2015; 10(5): e0122408.

13. Bolotov IN, Makhrov AA, Gofarov MY, Aksenova OV, Aspholm PE, Bespalaya YV, Kabakov MB, Kolosova YS, Kondakov AV, Ofenböck T, Ostrovsky AN, Popov IY, von Proschwitz T, Rudzīte M, Rudzītis M, Sokolova SE, Valovirta I, Vikhrev IV, Vinarski MV, Zotin AA. Climate warming as a possible trigger of keystone mussel population decline in oligotrophic rivers at the continental scale. Scientific Reports. 2018; 8: 35. doi:10.1038/s41598-017-18873-y

14. Klishko OK, Lopes-Lima M, Froufe E, Bogan AE. Taxonomy, morphometry and distribution of *Crictaria plicata* (Bivalvia, Unionidae) from the Russian Far East. ZooKeys. 2016; 580: 13–27.

15. Klishko OK, Lopes-Lima M, Froufe E, Bogan AE, Vasilieva L,., Yanovich L. Taxonomic reassessment of the *Unio* species (Bivalvia: Unionidae) from Russia and Ukraine based upon morphological and molecular data. Zootaxa. 2017; 4286(1): 93–112.

16. Klishko OK, Lopes-Lima M, Froufe E, Bogan AE, Abakumova V. Unravelling the systematics of *Nodularia* (Bivalvia, Unionidae) species from eastern Russia. Syst Biodivers. 2018; 16(3): 287–301. http://dx.doi.org/10.1080/14772000.2017.1383527.

17. Lopes-Lima M, Burlakova LE, Karatayev AY, Mehler K, Seddon M, Sousa R. Conservation of freshwater bivalves at the global scale: diversity, threats and research needs. Hydrobiologia. 2018; 810: 1–14.

18. Ferreira-Rodríguez N, Akiyama BY, Aksenova O, Araujo R, Barnhart C, Bespalaya Y, Bogan A, Bolotov IN, Budha PB, Clavijo C, Clearwater SJ, Darrigran G, Do VT, Douda K, Froufe E, Graf D, Gumpinger C, Humphrey CL, Johnson NA, Klishko O, Klunzinger MW, Kovitvadhi S, Kovitvadhi U, Lajtner J, Lennart H, Lopes-Lima M, Moorkens EA, Nagayama S, Nagel KO, Nakano M, Negishi J, Ondina P, Oulasvirta P, Pfeiffer P, Prié V, Riccardi N, Rudzīte M, Seddon M, Sheldon F, Sousa R, Strayer DL, Takeuchi M, Taskinen J, Teixeira A, Tiemann J, Urbanska M, Varandas S, Vinarski M, Wicklow BJ, Zajac T, Vaughn CC. Research priorities for freshwater mussel conservation assessment. Biol Conserv. 2019; 231: 77–87.

19. Mitchell J. Prehistoric Molluscan Faunas of the Yazoo River, Mississippi, USA: Archaeological Perspectives for Modern Conservation. Environ Archaeol. 2017; doi: 10.1080/14614103.2017.1288886

20. Joordens JCA, d’Errico F, Wesselingh F (and other 18 authors). *Homo erectus* at Trinil on Java used shells for tool production and engraving. Nature 2015; 518: 228–231.

21. Joordens JCA, Wesselingh F, de Vos J, Vonhof HB, Kroon D. Relevance of aquatic environments for hominins: a case study from Trinil (Java, Indonesia). J Hum Evol. 2009; 57(6): 656–671.

22. Lutaenko KA, Artemieva NG. Mollusks from the shell-midden of the Telyakovskogo 2 site in southern Primorye (Yankovskaya culture), their paleoecology and role in paleoeconomy. Bull Russ Far East Malacol Soc. 2017; 21(1/2): 61–128. (In Russian).

23. Lazăr C, Mărgărit M, Radu V. Evidence for the production and use of *Lithoglyphus naticoides* beads in Europe during the Holocene: The case of Sultana-Malu Rosu site (Romania). Quatern Int. 2018; 472(A): 84–96.

24. Preobrazhenskiy VC. About vertical zoning in the intermountain trough. Proceedings of the Academy of Sciences USSR. Geography series. 1958; 3: 151–168. (In Russian).

25. Zhukov VM. General features of climate. Types of territory and natural regionalization of the Chita oblast’. In: Popov SD, Preobrazhenskiy VC, editors. Proceedings of the Academy of Sciences USSR; 1961. 158 p. (In Russian).

26. Yendrikhinsky AS. The problems of paleolimnology and climatic stratigraphy of the Late Cenozoic period. Late Cenozoic history of the lakes in the USSR. In: Adamenko OM, Galazy GI, Belova VA., editors. Novosibirsk: Nauka, Siberian Branch; 1982. pp. 173–180. (In Russian).

27. Karasev VV. 2002. The Cenozoic of Transbaikalye. Chita: Chita regional publishing house; 2002. (In Russian).

28. Enikeev FI, Potemkina VI, Staryshko VE. Stratigraphy and evolution of climate and vegetation of late Cenozoic of north Transbaikalia. Novosibirsk: Geo; 2013. (In Russian).

29. Reshetova SA. Subrecent pollen spectra of the southern part of Transbaikalia as a methodlogical basis for the reconstructions of paleovegetation. Geografiya i prirodnyye resursy. 2018a; 1: 186–196. (In Russian).

30. Reshetova SA. Reconstruction of vegetation of the Chita-Ingoda depression (Transbaikalia) in the Late Holocene. Geosphere Research. 2018b; 4: 56–63.

31. Reshetova SA., Bezrukova EV. The vegetation and climate of Transbaikalia in the late glacial and holocene (by palynological data). Chita: ZabSU; 2018. (In Russian).

32. Okladnikov AP. Ancient Transbaikalia (cultural and historical essay). Life and art of the Russian population of the Eastern Siberia. In 2 parts. P. 2. Transbaikalia. In: Makovetski VI, Maslova GR, editors. Novosibirsk: Nauka, Siberian Branch; 1975. pp. 6–20. (In Russian).

33. Okladnikov AP, Larichev VE. Archaelogical investigations in the Amur Basin in 1954. In: Bolotin DP, Zabiyako AP, editors. Blagoveshchensk: AmurSU; 1999. pp. 4–29. (In Russian).

34. Kovychev EV, Kharinskii AV, Kradin NN. Shiwei’s fortress Proezzhaya 1 on the Shilka River. Journal of Ancient Technology Laboratory. 2018; 14(4): 78–102. doi: 10.21285/2415-8739-2018-4-78-102. (In Russian).

35. Kulakov VS., editor. Atlas of Transbaikalsky kray. Archeological map. Chita; 2010. p. 47. (In Russian).

36. Klishko OK. *Dahurinaia transbaicalica* sp. n. (Bivalvia, Margaritifridae)–new species of pearl mussel from Transbaikalie with remarks on natural history of the Far Eastern nayads. Vestnik zoologii. 2008; 42(4): 291–302.

37. Bolotov IN, Kondakov AV, Konopleva ES, Vikhrev IV, Aksenova OV, Aksenov AS, Bespalaya YV, Borovskoy AV, Danilov PP, Dvoriankin GA, Gofarov MY, Kabakov MV, Klishko OK, Kolosova YS, Lyubas AA, Novoselov AP, Palatov DM, Savvinov GN, Solomonov NM, Spitsin VM, Sokolova SE, Tomilova AA, Froufe E, Bogan AE, Lopes-Lima M, Makhrov AA, Vinarski MV. Integrative taxonomy, biogeography and conservation of freshwater mussels (Unionidae) in Russia. Scientific Reports. 2020; 10: 3072. https://doi.org/10.1038/s41598-020-59867-7.

38. Lopes-Lima M, Hattori A, Kondo T, Lee JH, Kim SK, Shirai A, Hayashi H, Usui T, Sakuma K, Toriya T, Sunamura Y, Ishikawa H, Hoshino N, Kusano Y, Kumaki H, Utsugi Y, Yabe S, Yoshinari Y, Hiruma H, Tanaka A, Sao K, Ueda T, Sano I, Miyazaki JI, Gonçalves DV, Klishko OK, Konopleva ES, Vikhrev IV, Kondakov AV, Gofarov MY, Bolotov IB, Sayenko EM, Soroka M, Zieritz A, Bogan AE, Froufe E. Freshwater mussels (Bivalvia: Unionidae) from the Rising Sun (Far East Asia): Phylogeny, systematics, and distribution. Molec Phyl Evol. 2020; 146: 106755. https://doi.org/10.1016/j.ympex.2020.106755.

39. Starobogatov YI, Prozorova LA, Bogatov VV, Sayenko EM. Mollusca. In: Tsalolikhin SY, editor. Key to freshwater invertebrates of Russia and adjacent lands. Vol. 6. Molluscs, Polychaetes, nemerteans. Saint Petersburg: Nauka; 2004. pp. 6–402. (In Russian).

40. Wesselingh FP, Neubauer TA, Anistratenko VV, Vinarski MV, Yanina T, ter Poorten JJ, Kijashko P, Albrecht C, Anistratenko OY, D’Hont A, Frolov P, Gándara AM, Gittenberger A, Gogaladze A, Karpinsky M, Lattuada M, Popa L, Sands AF, Van de Velde S, Vandendorpe J, Wilke T. Mollusc species from the Pontocaspian region–an expert opinion list. ZooKeys. 2019; 827: 31–124.

41. Kirilov II, Dyatchina NG. Duroy, the old relict of archeology. In: Geniatulin RF, editor. Encyclopedia of Transbaikalia. Novosibirsk: Nauka; 2004. Vol. 2. p. 328. (In Russian).

42. Evseev GA. Bivalves of the South Kuril shallow waters and their habitats. Bull Russ Far East Malacol Soc. 2000; 4: 30–51.

43. Lutaenko КA. Bivalve molluscs of the Amursky Bay (Sea of Japan/East Sea) and adjacent areas. Part 1. Families Nuculidae–Cardiidae. Bull Russ Far East Malacol Soc. 2002; 6: 5–60.

44. Sayenko EM, Prokopets SD, Lutaenko KA. Molluscs from medieval Bohai settlement Nikolaevskoe I (Primorye, Russian Far East): paleecological and archeozoological significance. Ruthenica: The Russian Malacological Journal. 2015; 25(2): 51–67.

45. Yaremchuk OA. Zorgol archeological culture. In: Geniatulin RF, editor. Encyclopedia of Transbaikalia. Novosibirsk: Nauka; 2004. Vol. 2. pp. 414–415. (In Russian).

46. Prozorova LA, Sitnikova TY, Zasypkina MO, Matafonov PV, Dulmaa A. Freshwater gastropods (Gastropoda) of the Baikal Lake basin and adjacent areas. In: Timoshkin OA, editor. Index of animal species inhabiting lake Baikal and its catchment area. Vol. 2. Basins and channels of the south of East Siberia and Mongolia. Novosibirsk: Nauka Publishers. 2009; 1: 170–188. (In Russian).

47. Volkov IA, Volkova MC. Late glacial and Holocene history of the lakes of the southern part of the Western Siberian Plain according to the geological data. Late Cenozoic history of the lakes in the USSR. In: Adamenko OM, Galazy GI, editors. Novosibirsk: Nauka, Siberian Branch; 1982. pp. 101–108. (In Russian).

48. Popova SM. Cenozoic continental malacofauna of south Siberia and adjacent lands. Moscow: Nauka; 1981. (In Russian).

49. Kulchitski AA, Misharina VA, Popova SM, Adamenko RS. To biostratigraphy and paleogeography of the lower and middle Pleistocene of the Kirenga River Basin. In: Adamenko OM, editor. Materials on biostratigraphy and paleogeography of the Eastern Siberia. Moscow: Nauka; 1975. 158p. (In Russian).

50. Vinarski MV, Kantor YI. Analytical catalogue of fresh and brackish water molluscs of Russia and adjacent countries. Moscow: A.N. Severtsov Institute of Ecology and Evolution of Russian Academy of Sciences; 2016.

51. Glöer P. The freshwater gastropods of the West-Palaearctis. Volume 1. Fresh- and brakish waters except spring and subterranean snails. Identification key, anatomy, ecology, distribution. 2019; Published by the author.

52. Dybowski B. Bemerkungen und Zusatze zu der Arbeit von Dr. W. Dybowski “Mollusken der Uferregion des Baikalsees.” Ezhegodnik Zoologicheskogo Muzeya Imperatorskoy Akademii Nauk. 1913; 17: 165–218.

53. Dybowski W. Mollusken aus der Uferregion des Baikalsees. Ezhegodnik Zoologicheskogo Muzeya Imperatorskoi Akademii Nauk. 1913; 17: 123–143.

54. Økland J. Lakes and snails: environment and Gastropoda in 1,500 Norwegian lakes, ponds and rivers. Oegstgeest: Universal Book Services/W. Backhuys; 1990.

55. Vinarski MV, Karimov AV. Geographic variation of *Planorbis planorbis* shells in the waterbodies of Western Siberia (Gastropoda: Pulmonata: Planorbidae). Mollusca (Dresden). 2008; 26(2): 195–206.

56. Khokhutkin IM, Vinarski MV. Molluscs of the Urals and the adjacent areas. The families Acroloxidae, Physidae, Planorbidae (Gastropoda, Pulmonata, Lymnaeiformes). Fasc. 2. Yekaterinburg: Goshchitskiy Publishers. 2013; 184 p. (In Russian).

57. Enikeev FI, Staryshko VE. Glacial morphogenesis and placer formation of the Eastern Transbaikalia. Chita: ZabSU; 2009. (In Russian).

58. Enikeev FI. Glacial loading basins of Eastern Transbaikalye: new look on the paleogeography of the Pleistocene. Geology and Geophysics. 2018; 59(9): 1384–1396.

59. Zhadin VI. Family Unionidae. Fauna of the USSR. Mollusks. 1938; 4(1): 1–170. (In Russian).

60. Konstantinov MV. The Stone Age of the eastern region of Baikalian Asia. Ulan-Ude: Buryat Scientific Center; Siberian Branch of the Russian Academy of Sciences; Chita State Pedagogical Institute; 1994. 179 p. (In Russian).

